# A single *de novo* substitution in SARS-CoV-2 spike informs enhanced adherence to human ACE2

**DOI:** 10.1101/2021.07.16.452441

**Authors:** Elena Erausquin, Jacinto López-Sagaseta

**Affiliations:** Unit of Protein Crystallography and Structural Immunology, Navarrabiomed, 31008, Navarra, Spain; Public University of Navarra (UPNA), Pamplona, 31008, Navarra, Spain; Navarra Hospital Complex (CHN), Pamplona, 31008, Navarra, Spain

## Abstract

SARS-CoV-2 initiates colonization of host cells by binding to cell membrane ACE2 receptor. This binding is mediated by the viral spike receptor binding domain (RBD). The COVID-19 pandemic has brought devastating consequences at a clinical, social and economical levels. Therefore, anticipation of potential novel SARS-causing species or SARS-CoV-2 variants with enhanced binding to ACE2 is key in the prevention of future threats to come. We have characterized a *de novo* single substitution, Q498Y, in SARS-CoV-2 RBD that confers stronger adherence to ACE2. While the SARS-CoV-2 β variant, which includes three simultaneous amino acid replacements, induces a 4-fold stronger affinity, a single Q498Y substitution results in 2.5-fold tighter binding, compared to the Wuhan-Hu-1 SARS-CoV-2 2019 strain. Additionally, we crystallized RBD_Q498Y_ complexed with ACE2 and provide here the structural basis for this enhanced affinity. These studies inform a rationale for prevention of potential SARS-causing viruses to come.

## Introduction

As the earlier severe acute respiratory syndrome coronavirus (SARS-CoV) (*1*) and the Middle East Respiratory Syndrome (MERS) coronavirus (*2*, *3*), SARS-CoV-2 (*4*) belongs to the genus betacoronavirus, and is the causative agent of the Coronavirus disease 2019 (Covid19) pandemic that stroke mankind in late 2019.

Covid19 is currently an ongoing threat counteracted by means of massive campaigns of vaccination in humans. While vaccination routines associate with a reduction in severity and clinical outcome, new SARS-CoV-2 variants emerge and pose new challenges for which there is no currently a universal preventive measurement.

Since the first Covid19 outbreak in 2019, new SARS-CoV-2 variants are surfacing and have triggered new viral outbreaks (*5*, *6*). Some of these variants are characterized by mutations that affect the RBD binding site on ACE2. A variant bearing the N501Y was detected in the United Kingdom, referred as Variant of Concern 202012/01 or alpha variant, following the World Health Organization convention (*7*), and has been linked to a higher transmissibility (*8*). The 501.V2 variant, or beta variant, was originally identified in South Africa in December 2020. While the prevalence of this variant was higher among the youths (*9*), it correlates with a more severe clinical condition. Further, this variant appears to disseminate at a higher rate, compared to previously identified SARS-CoV-2 variants. SARS-CoV-2 β holds three amino acid substitutions in the RBD. These are N501Y, K417N and E484K, which appear to enhance the binding strength of the spike to ACE2 (*10*).

The delta variant was originally identified in India, is already predominant in many countries and includes L452R and T478K substitutions in the RBD. The delta plus variant includes an extra amino acid replacement, K417N (*11*).

Additionally, increased evidences point to transmissions to humans due to zoonosis. Zoonosis poses further challenges as novel infectious pathogens can be spread from animals to humans. For instance, from 2017 to 2020, five zoonotic avian-related transmissions to humans have been reported (*12*). Since 2003, three coronaviruses have brought devastating consequences to humans at clinical, economic and social levels. Investigation and anticipation are therefore critical in the prevention of novel pathogens to come with potential to infect humans, in the form of either novel viruses or variants of currently circulating germs.

SARS-CoV-2 initiates contacts with the host through its viral envelope-anchored spike (S) protein, which binds with very high affinity to the Angiotensin-converting enzyme 2 (ACE2) on the surface of human cells (*13*). Many structural studies have evinced with great detail the structural and molecular blueprint of this binding, which involves a nourished array of both polar and non-polar interactions between both the spike RBD and ACE2 (*14*–*16*). The binding interface covers a molecular area of 850 Å2. Eighteen residues on the spike RBD establish direct interactions with ACE2 through bonds of different nature, including hydrogen bonds, Van der Waals forces and salt bridges.

Following the aforementioned need to inform potential novel threats, we sought to identify amino acid substitutions that might enhance the adherence for ACE2. We performed an *in silico* mapping of the RBD binding interface and identified a single substitution, Q498Y, that was predicted to enhance binding to ACE2. We provide evidences for this increased adherence using biolayer interferometry approaches. In addition, we report structural grounds that support the molecular mechanism underlying this increased affinity. Genetic mutations leading to Q>Y replacements are not within the most likely variations to occur in nature. Nevertheless, the clinical records collect several isolates of Q>Y replacements in humans infected with SARS-CoV-2.

These studies inform a *de novo* substitution in SARS-CoV-2 that results in a tighter binding to ACE2 and which might lead to a compromised clinical setting.

## Results

### *In silico* prediction of RBD mutations that enhance binding to ACE2

Using a computational pipeline for high-throughput *in silico* mutagenesis, we explored substitutions in the RBD wild type (RBD_WT_)-human ACE2 (ACE2) interface leading to a predicted increase in affinity by means of a reduced free binding energy score.

The mutational blueprint calculated for the RBD-ACE2 interface (Fig. 1), showed a heterogeneous plot whereby position 475 in RBD, corresponding to Alanine, gathers the most favorable substitutions in terms of affinity. In contrast, Tyr489 is, apparently, the less favorable residue to enhance binding to ACE2 upon substitution. Looking at the chemical nature of the replacement amino acid, substitutions to Tyr and Pro result in the most and less beneficial variations, respectively. Here, we focused on the substitution with the greatest score, Q498Y, which produced a predicted affinity change (ΔΔG) of 2.0 kcal/mol, to experimentally verify its impact on the binding affinity of RBD for ACE2. Gln498 is located within the RBD-ACE2 binding interface, particularly in the RBD region near Tyr41 and Gln42 in ACE2 N-terminal α-helix, with which establishes polar and VDW contacts (Fig. 1B). More specifically, there is a putative H-bond with Gln42 side chain at a distance of 3.4 Å. VDW interactions are more populated and participate in binding both Tyr41 and Gln42.

**Figure 1.**
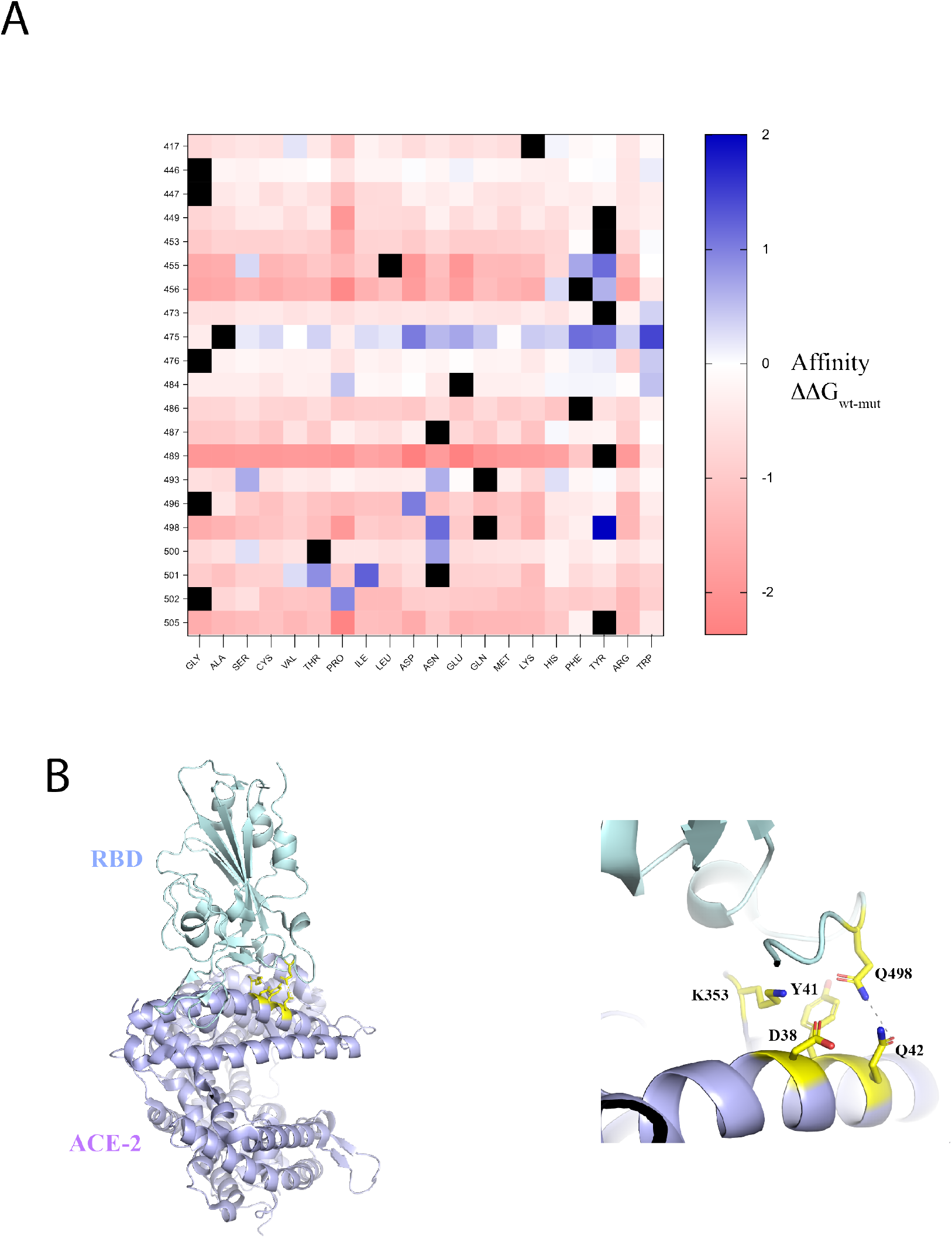
Mutational blueprint of RBD-ACE2 interface. (A) Heatmap plotting changes in predicted binding affinity of RBD_WT_ to ACE2. The RBD binding interface was *in silico* (*17*) screened for amino acid substitutions leading to a predicted enhanced affinity for ACE2. The working template was obtained from the X-ray coordinates deposited in the Protein Data Bank under the accession code 6M0J. (B), Overall cartoon representation of the RBD_WT_-ACE2 crystal structure (left panel) with position 498 in RBD_WT_ and surronding residues highlighted in yellow color. The right panel shows a closer image of Gln498 in RBD_WT_. The dashed line represents a likely (3.4 Å) hydrogen bond between RBD_WT_ Gln498 and ACE2 Gln42.

### A single Q>Y substitution in position 498 of RBD confers greater binding to ACE2

In order to investigate the Q498Y mutation, we produced recombinant human ACE2, RBD_WT_ RBD_Q498Y_ and RBD β proteins in sf9 cells. Purified proteins were used to perform biolayer interferometry assays for an accurate characterization of the binding affinities. We first captured ACE2 on the surface of a nickel sensor through a 12xHis c-terminal tag. Increasing concentrations of RBD_wt_, RBD_Q498Y_ or RBD_β_ were then loaded and the association and dissociation curves were monitored in real time (Fig. 2). The RBD_β_ version served as a reference for a naturally-ocurring SARS-CoV-2 variant (Fig. S1).

**Figure 2.**
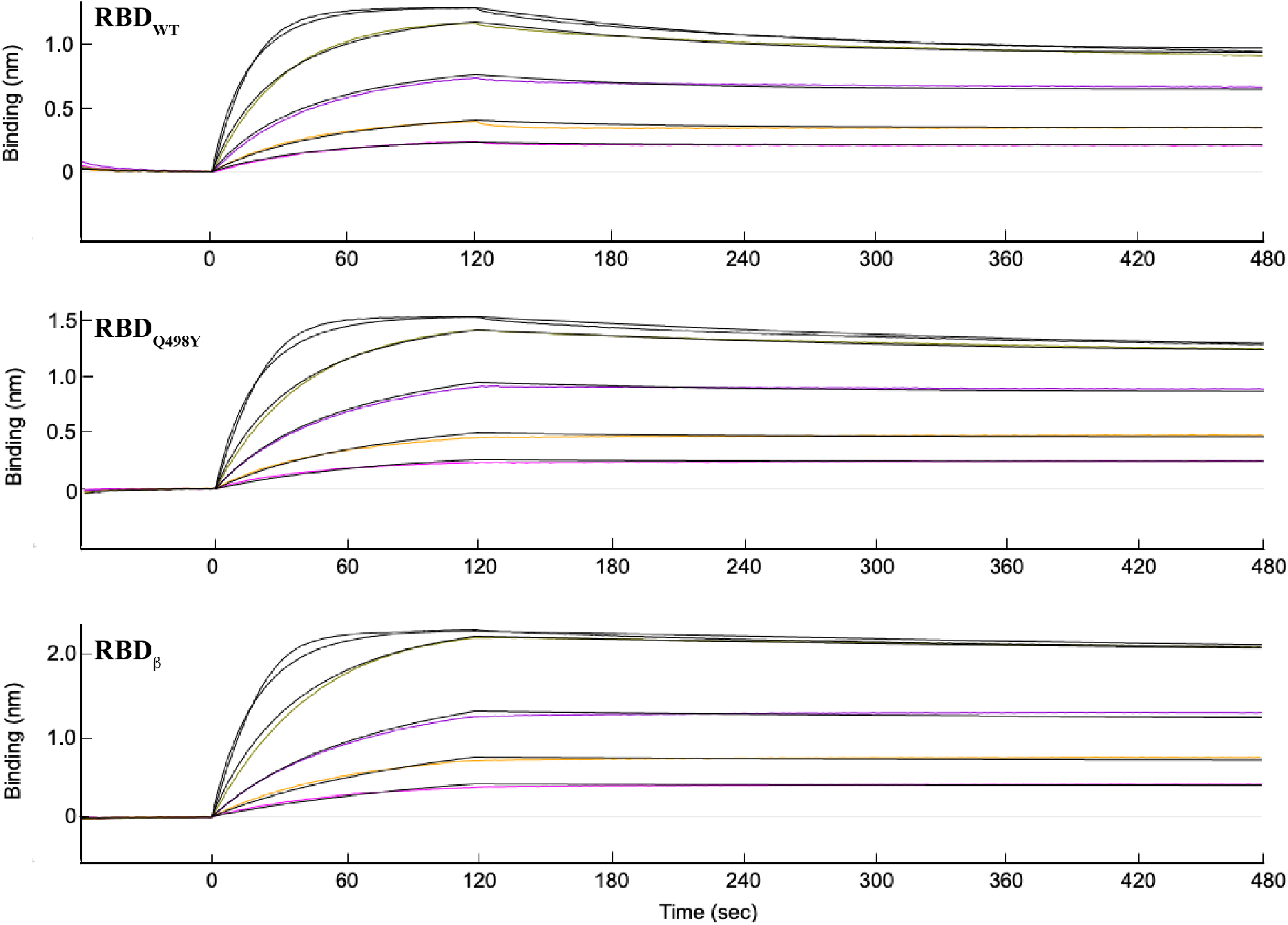
RBD_Q498Y_-ACE2 binding kinetics determined by biolayer interferometry (BLI). (A) ACE2 was captured on the surface of a NiNTA sensor and pulsed for 2 minutes with increasing concentrations of RBD_WT_, RBD_Q498Y_ or RBD_β_ in running buffer (20 mM Hepes pH 7.4, 150 mM NaCl). The association and dissociation data were monitored and fitted to a 1:1 Langmuir model for calculation of binding kinetics constants (*k_on_*, *k_off_* and *K*_D_) and assessment of fitting goodness (R^2^). (B) Table summarizing the calculated parameters of RBD_WT_, RBD_Q498Y_ or RBD_β_ binding to ACE2.

We could record a remarkably stable binding with all three RBD forms tested, where a tendency for a modestly slower dissociation rate could be noticed for the Q498Y and β forms of the RBD (Fig. 2). Varied response magnitudes could be likewise detected, with RBD_WT_ and RBD_β_ producing the lowest and highest values, respectively. A more accurate comparison was obtained by determining the kinetic constants of each interaction. The data could be fitted to a 1:1 Langmuir model with coefficients of determination R^2^ above 0.99 in all cases. All three RBD forms bound ACE2 with very high affinity and K_D_ values near subnanomolar range (Table I). However, among the three RBD forms tested, RBD_Q498Y_ showed the fastest association rate, yet with a *k_on_* value very similar to those of RBD_WT_ and RBD_β_ (Table I). The main difference between RBD_Q498Y_ and RBD_WT_ fell in the dissociation rate, where RBD_Q498Y_ showed a ~ 2.5-fold lower *k_off_* compared to that of RBD_WT_. Thus, RBD_Q498Y_ resulted in an overall ~ 2.5-fold tighter binding. The RBD_β_ variant represented the most stable binder with a 2.8×10^−4^ s^−1^ *k_off_* value, i.e., ~2 and ~4.5-fold slower dissociation rate than RBD_Q498Y_ and RBD_WT_, respectively. Nevertheless, and despite the three amino acid substitutions, the overall binding affinity of the RBD_β_ form is only 1.6 times stronger than that of RBD_Q498Y_ (Fig. 1, Table I). This results in a relative change in affinity of 1.4-fold per amino acid replaced, while a single Q498Y replacement confers a 2.5-fold increased in binding affinity.

**Table I.** Kinetic constants determined by biolayer interferometry. Shown are the association and dissociation constant parameters, *k_on_* and k_off_, respectively, as well as the overall K_D_ (*k_off_/k_on_*) value. The gooodness of the binding data fittings to a 1:1 Langumir model is represented by the R^2^ or coefficient of determination.

| | $k_{on} \times 10^5 (M^{-1} s^{-1})$ | $k_{off} \times 10^{-4} (s^{-1})$ | $K_D (nM)$ | $R^2$ |
| --- | --- | --- | --- | --- |
| RBD wt | $1.3 \pm 0.3$ | $13.0 \pm 6.9$ | $9.4 \pm 2.8$ | 0.9980 |
| RBD Q498Y | $1.5 \pm 0.2$ | $5.2 \pm 3.1$ | $3.7 \pm 2.6$ | 0.9982 |
| RBDβ | $1.2 \pm 0.1$ | $2.8 \pm 0.5$ | $2.3 \pm 0.6$ | 0.9975 |

To further confirm this trend, we assembled an additional but indirect BLI approach whereby RBD_wt_ was covalently immobilized on the surface of an AR2G sensor. ACE2 at a concentration of 400 nM was then pulsed and the binding curves were compared with similar concentrations of ACE2 but previously incubated with stoichiometric amounts of RBD_WT_, RBD_Q498Y_ or RBD_β_ (Fig. 3), *i.e.,* we set out to investigate the binding strenght to ACE2 in solution. This alternative strategy allowed us to confirm a stronger binding between ACE2 and RBD_Q498Y_, when compared with RBD_WT_, as we could notice a stronger competition in the association of ACE2 to surface RBD_WT_. As in the direct assay, the RBD_β_ variant showed the tightest binding to ACE2, as mirrored in the strongest inhibition of ACE2 binding to surface-RBD_WT_ (Fig. 3).

**Figure 3.**
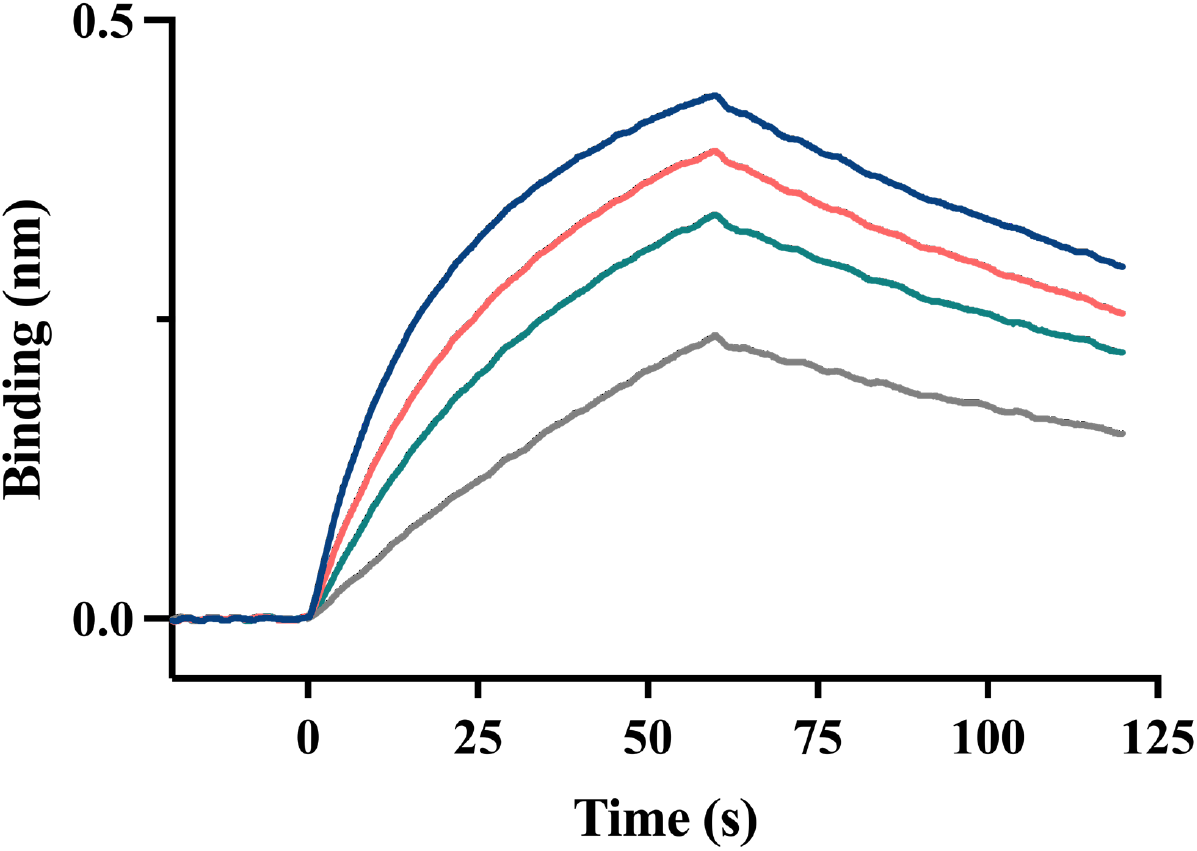
Indirect assesment of Q498Y subsitution in binding affinity to ACE2. A BLI competition assay was setup whereby RBD_WT_ was covalently captured on the surface of an amine-reactive sensor. ACE2 at a concentration of 400 nM (blue color line), free or preincubated in solution with 400 nM RBD_WT_ (orange line), RBD_Q498Y_ (teal line) or RBD_β_ (grey line) was then loaded over the RBD_WT_ surface and the binding profile was monitored.

### Structural bases for RBD_Q498Y_ enhanced affinity

To explore the structural basis for this enhanced affinity, we prepared a complex between RBD_Q498Y_ and ACE2 (Fig. 4). Incubation of RBD_Q498Y_ with ACE2 showed a sharp shift in the retention time of the protein through size exclusion column chromatography.

**Figure 4.**
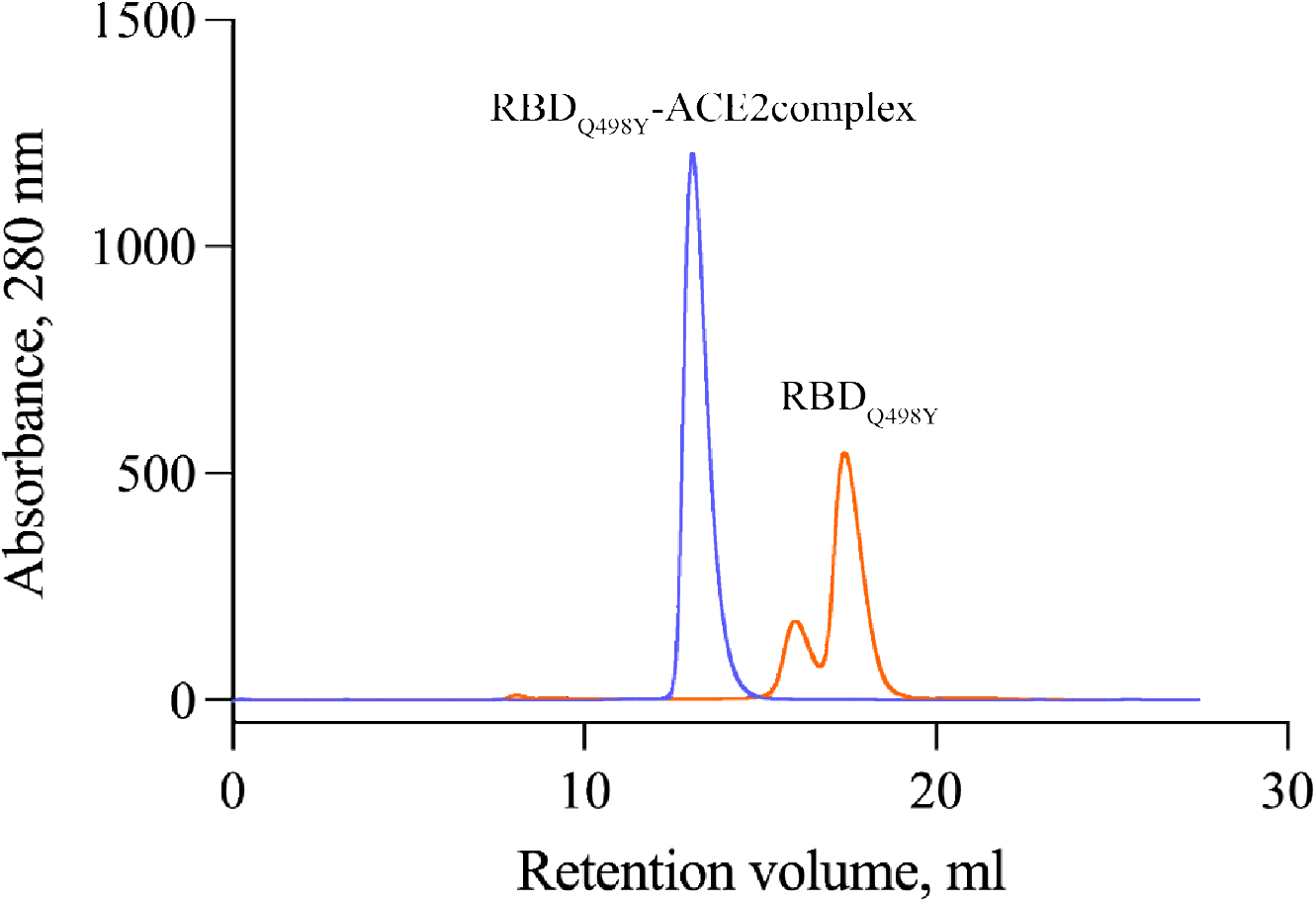
Formation of the RBD_Q498Y_-ACE2 complex. The elution profile of RBD_Q498Y_ alone (orange trace) or complexed with ACE2 (blue trace) was monitored to track a proper formation of the complex through size exclusion chromography.

We then screened over 750 crystallization conditions to obtain crystals of RBD_Q498Y_-complexed with ACE2. Crystals of the RBD_Q498Y_-ACE2 complex appeared in with 0.1 M sodium phosphate pH 6.5 and 12% PEG 8000. A full dataset to 3.3 Å was collected and processed in P212121 space group. Initial phases were calculated with the molecular replacement method, and yield two RBD_Q498Y_-ACE2 complexes per assymetric unit. The overall docking mode is almost identical to that of the complex with RBD_WT_ (Fig. 5A). The binding footprint is preserved and shows an elongated interface where RBD_Q498Y_ interacts primarily with ACE2 N-terminal α-helix (Fig. 5 and Table S2). The buried surface area (BSA) of the interaction covers 820 and 837 Å^2^ in each complex structure in the assymmetric unit. As for the complex with RBD_WT_, a strong electron density signal is also found for a Zn^2+^ ion, which is coordinated by His374, His378, Glu375 and Glu402 in ACE2. The electron density in position 498 allowed us identify a preserved backbone structure, with Tyr498 side chain projected towards the RBD. More specifically, Tyr498 side chain occupies a pocket in the RBD-ACE2 in a manner similar to that of Gln98 in the complex structure with the wild type counterpart (Fig. 5B-C). However, this structural arrangement and the bulkier side chain of tyrosine in RBD_Q498Y_ promotes differences with respect to glutamine in RBD_WT_.

**Figure 5.**
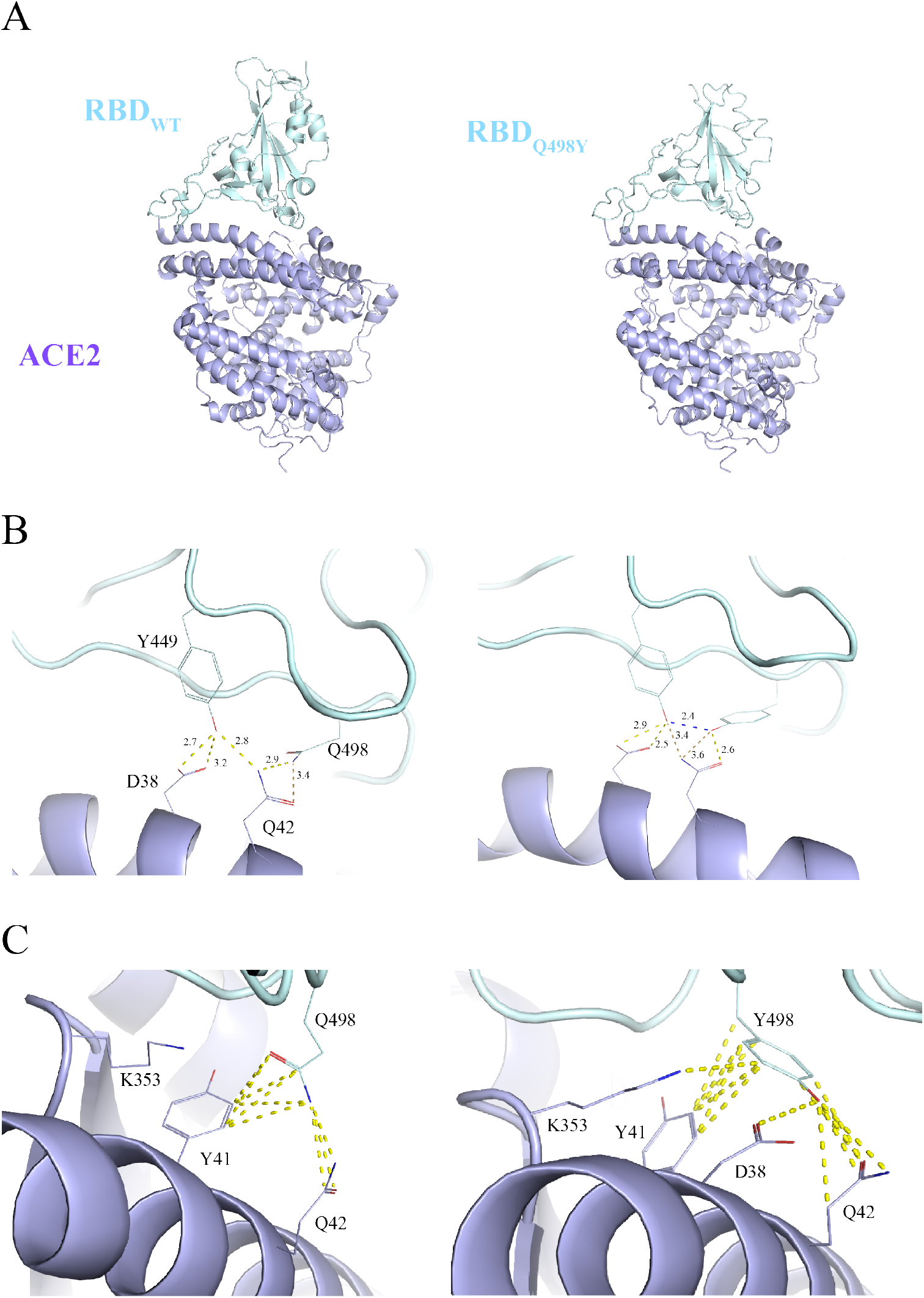
Structural features of the RBD_Q498Y_-ACE2 complex. (A), Comparison of the overall complex structures of ACE2 complexed with RBD_WT_ (image prepared with the atomic coordinates deposited in the Protein Data Bank, accession code 6M0J) or RBD_Q498Y_, as indicated in the figure. (B), Intermolecular interactions (dashed lines) mediated by hydrogen bonding. Interacting residues are depicted as sticks. H-bond distances are shown. (C), Intermolecular, non-polar interactions are indicated with dashed lines.

In the complex structure with the RBD_WT_ structure, Gln498 contacts Gln42 in ACE2 through a single 3.4 Å H-bond. On the contrary, the presence of the bulky Tyr498 side chain promotes a minor alteration of Gln42 from its position in the complex with RBD_WT_ to accommodate the long tyrosine side chain (Fig. 6A). This arrangement favors, on one side, a shorter (2.6 Å) H-bond between Tyr498 side chain oxygen and Gln42 side chain oxygen in ACE2, and an additional putative 3.6 Å H-bond with Gln side chain nitrogen (Fig. 5 and Table S2). Moreover, Tyr498 locks Tyr449 through a 2.4 Å-side chain-side chain H-bond, which consequently brings it closer to Asp38 in ACE2. Following this arrangement, the Tyr449-Asp38 H-bond distances are 2.9 Å and 2.5 Å with Asp33 side chain oxygens, vs 2.7 Å and 3.2 Å in the complex with RBD_WT_. In addition, the number of non-polar interactions is higher in the case of the RBD_Q498Y_ complex (Fig. 5C). These additional contacts are observed with Asp38, Gln 42 and Lys353. Altogether, the presence of a tyrosine instead of glutamine supports a more populated network of interactions that helps stabilize the RBD-ACE2 complex.

**Figure 6.**
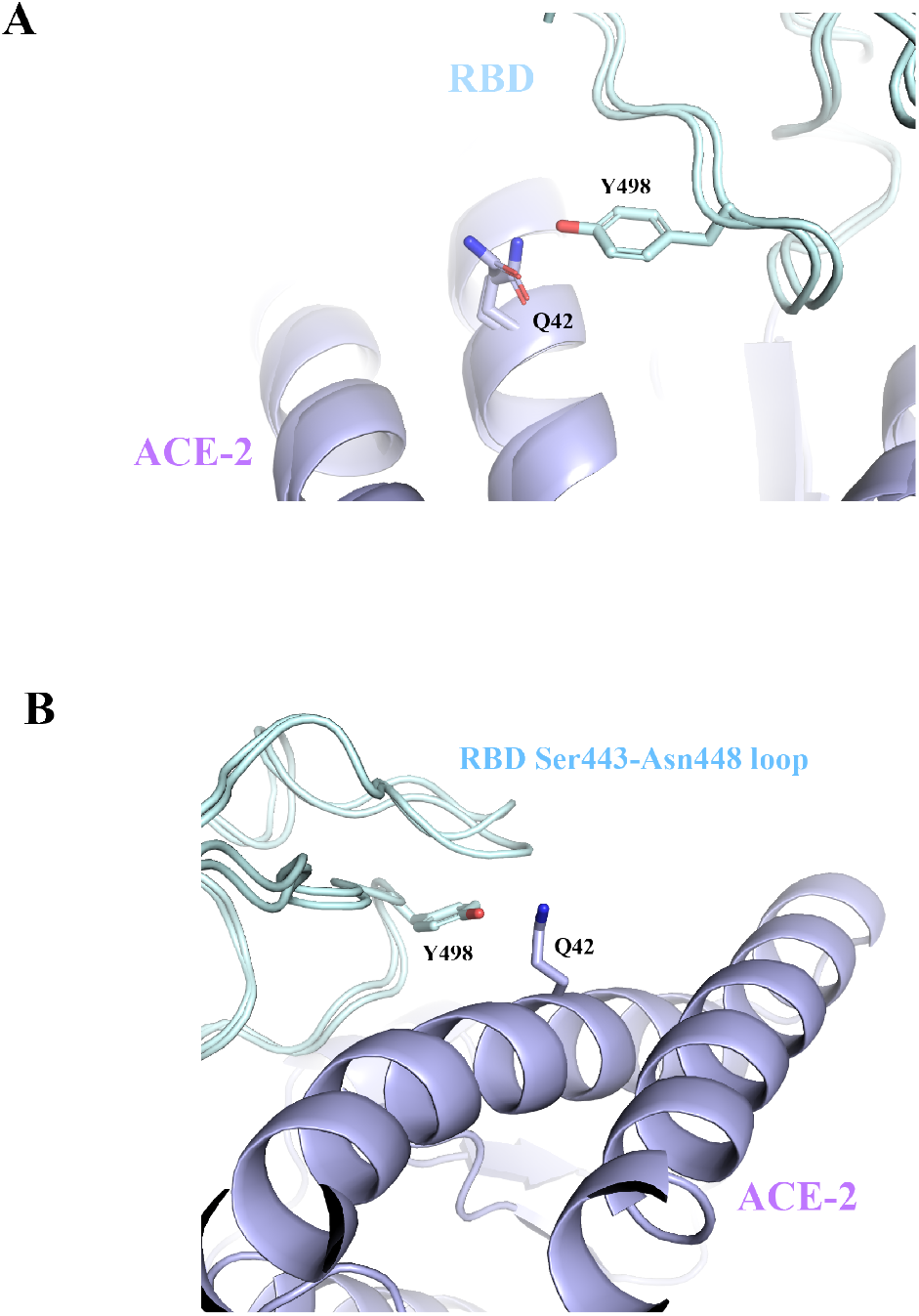
Conformational changes induced by Q498Y. (A), Cartoon representation showing Gln42 rotamer shift (highlighted with sticks) in the presence of RBD Tyr498. To appreciate the side chain movement, Q42 in the complex with RBD_WT_ is also represented as sticks. (B), the structural conformation of the Ser443-Asn448 loop in RBD_Q498Y_ is compared with that of the complex with RBD_WT_. An upwards displacement can be observed in the presence of Tyr498 in RBD_Q498Y_.

### Conformational changes induced by Q498Y substitution

Interestingly, we observed conformational changes induced by the Q498Y substitution. First, Gln42 shows a moderate displacement away from the interface (Fig. 6A). This displacement avoids steric hindrance and enables a suitable pocket for acommodation of Tyr498.

However, we could notice an additional conformational change in a loop delimited between Ser443 and Asn448 (Fig 6B). Again, the bulkier volume of Tyr498 suggests a rearrangement of this loop, which shifts its backbone conformation upwards. Two consecutive glycine residues in the loop, Gly446-Gly447, likely favor plasticity in this region. Thus, the SARS-CoV-2 RBD features a flexible RBD structure with ability for adaptation to mutations and preserve or even enhance binding to the human ACE2.

### Low prevalence of Q>Y substitutions

Aware of the low likelihood of a Q>Y substitution to occur, we looked into the prevalence of Q>Y substitutions reported in isolates from individuals infected with SARS-COV-2 as of April 2021 and relative to the Wuhan-Hu-1 strain genome. Analysis of putative substitutions identified in the GISAID database yield a total of 185 mutation events leading to Q>Y substitutions (Fig. 7). The highest prevalence falls in ORF3a, with 150 events identified. The other 35 events are distributed across the diverse open reading frames in the SARS-CoV-2 genome, and include 10 Q>Y replacements in the spike protein. However, no Q498Y substitution has yet been reported.

**Figure 7.**
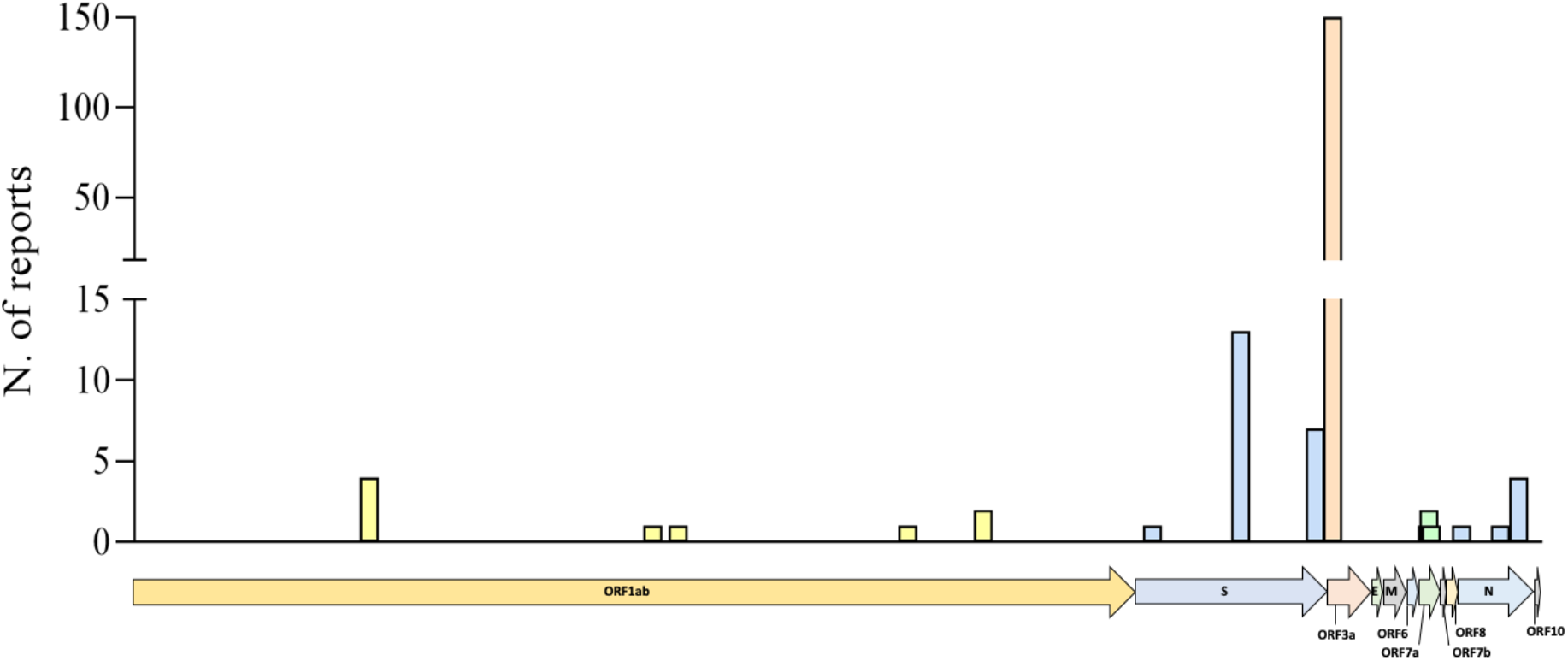
Mutational incidence for Q>Y substitutions in circulating SARS-CoV-2. Shown is the distribution and frequency (number of reports), along the SARS-CoV-2 genome, of the different Q>Y substitutions reported as of April 6th 2021 in the GISAID Database. Q>Y replacements were identified through CoV-GLUE tracking tool for SARS-CoV-2 genomic variation. All replacements are relative to the Wuhan-Hu-1 SARS-CoV-2 genome isolate.

## Discussion

In the last twenty years, three coronaviruses have posed devastating consequences for mankind. SARS-CoV emerged in 2002 and infections in humans was reported to have a zoonotic cause. SARS-CoV caused > 8000 infections and > 775 deaths across the world (*1*). The Middle-East respiratory syndrome (MERS) coronavirus surface later in 2012, when the first infection in humans was reported, being camels apparently the natural reservoir (*3*). In humans, the records show ~ 2500 infections in humans and > 850 deaths. MERS still poses a health threath in Middle East. In late 2019, a third coronavirus, SARS-CoV-2 (*4*), with devastating consequences has caused a pandemic with > 181 million infections in humans and 3,937,437 deaths as of 30^th^ June 2021. Further, SARS-CoV-2 has led to new variants that associate with new outbreaks, and consequently, adding severity to the overall impact in human population since the first cases were noticed in 2019.

Whether new betacoronaviruses species or variants will appear and strike global health is unknown. For this reason, anticipation and gain of knowledge on how new forms of betacoronaviruses can interact with host cells is key to inform clinical tools in the prevention against new coronaviruses species and variants to come.

Here, we harnessed an *in silico* computational pipeline to map the SARS-CoV-2 RBD-human-ACE2 binding interface and searched for putative substitution in the RBD that could lead to an enhanced affinity for ACE2. Of all possible substitutions, Q498Y resulted in the highest score, with a predicted affinity change (ΔΔG) of 2.004 kcal/mol over its wild type counterpart, i.e., Q498Y was predicted to further stabilize the SARS-CoV-2 RBD-human-ACE2 interaction. In order to validate this prediction, we expressed recombinantly human ACE2, RBD_WT_, and RBD_Q498Y_, and compared the binding kinetic profiles using biolayer interferometry. Our binding studies showed an enhanced binding with RBD_Q498Y_, thus confirming the predicted *in silico* values. To gain understanding on the structural bases for this enhanced affinity, we grew crystals of the RBD_Q498Y_-ACE2 complex and solved the structure at a resolution of 3.3 Å. The electron density maps were of sufficient quality to depict the RBD_Q498Y_-ACE2 complex, which had an almost identical docking mode when compared with the complex with RBD_WT_ (*14*, *15*). Importantly, we could observe relevant changes in the region surronding position 498 in the RBD. While the overall docking interface is preserved, we noticed conformational rearrangements, apparently, due to the presence of the bulkier tyrosine residue, as opposed to glutamine. In particular, Gln42 in ACE2 shifts away from its position in the crystal structure complex with RBD_WT_, while a 6-residue loop near position 498 moderately alters its conformation thanks to two consecutive glycine residues. Both conformational changes avoid steric clashes and hindrance to accommodate Tyr498 side chain. Despite these structural rearrangements, Tyr498 provides a more populated network of polar and hydrophobic interactions with ACE2 that supports the findings through biolayer interferometry, and ultimately points to a tighter stabilization of the RBD-ACE2 complex.

Interestingly, the RBD of SARS-CoV, which originated a SARS epidemic in 2002, contains a tyrosine in the position equivalent to RBD498 in SARS-CoV-2 (*18*). A similar *in silico* evaluation on the impact of a reverse Y>Q replacement in binding affinity results in a predicted affinity change of −1.605 kcal/mol, *i.e.,* Y>Q substitution is less advantageous in terms of affinity at this particular position, which supports the contribution of Tyr498 to the overall affinity for ACE2.

From a genetic perspective, Q>Y replacements are not probabilistically favorable. Gln amino acids are translated as either CAA or CAG codons, while tyrosine residues are the result of UAU and UAC codons, *i.e.*, double mutations are required for a Q>Y replacement to occur. Still, the GISAID initiative has registered 185 Q>Y events that are distributed throughout the SARS-CoV-2 genome. The highest incidence falls in the open reading frame 3, which accounts for 150 reports, while the spike region, with 20, is the second region in the SARS-CoV-2 genome that concentrates the largest number of Q>Y substitutions.

Altogether, Q>Y replacements should not be disregarded despite their low likelihood, as any mutation leading to a more favorable viral evolution might pose a risk for humans as host targets of betacoronaviruses.

## Material and Methods

### In silico mutational screening

We performed a high-throughput *in silico* mutagenesis using structural information from the interface between RBD and ACE2. This information was collected from the atomic coordinates deposited in the Protein Data Bank under the accession code 6M0J and used to feed the mCSM-PPI2 pipeline (*17*). The method relies on the structure of a protein-protein template, the reference residue environment, the physicochemical properties of the replacement residue, the degree of evolutionary conservation and the thermodynamics of the protein-protein complex to predict the impact of such mutations. An amino acid substitution matrix focused on the wild type RBD binding site was generated and used to plot a heatmap to evaluate variations in protein-protein binding energy, determined as changes in Gibbs free energy (ΔΔG) upon single-residue mutagenesis.

### Cloning and generation of recombinant baculovirus

Codon-optimized gene sequences for human extracellular ACE2 containing a C-terminal 12xHis tag, and for SARS-CoV-2 wild type spike receptor binding domain (RBD) with a N-terminal TwinStrep tag were synthesized by BioBasic Inc. RBD beta variant RBD (RBD_β_) was synthesized by GeneUniversal, being position 334 the first N-terminal native amino acid, and which we referred as short RBD. Sequences were digested from delivery generic vectors using BamHI and NotI restriction enzymes (FastDigest) and cloned in the pAcGP67A transfer vector using Optizyme™ T4 DNA Ligase (Thermo Fisher Scientific). *E. coli* DH5α cells (Invitrogen) were transformed with the modified transfer vectors and before plasmid DNA extraction using the GeneJET Plasmid Miniprep Kit (Thermo Fisher Scientific) following manufacturer’s instructions. An N-terminal TwinStrep tag and 3C site containing version of ACE2 with no His tag was generated by PCR using ACE2 plasmid DNA as template. N-terminal TwinStrep-3C site short wild type RBD (RBD_WT_) and a similar construct including the Q498Y substitution (RBD_Q498Y_) were generated by PCR using specific primers. All new PCR products were further digested and cloned into pAcGP67A vectors as described before.

Once sequences were validated by Sanger sequencing, Sf9 insect cells (Gibco) were transduced with each transfer plasmid, BestBac 2.0 Δ v-cath/chiA Linearized Baculovirus DNA and Expres2 TR Transfection Reagent (Expression Systems) to produce the final recombinant baculovirus.

### Recombinant protein expression and purification

#### His-tagged protein expression and purification

Sf9 insect cells were infected with His-tagged ACE2 baculovirus, in a 1:2000 dilution, and left in agitation for 72 hours. Culture volume was collected, then supplemented with 40 mM HEPES, 300 mM NaCl and pH adjusted to 7.2. 40 mM imidazole, 5 mM MgCl_2_ and 0,5 mM NiSO_4_ were added to the sample. Recombinant protein was purified from the supernatant using HisGraviTrap (Cytiva) columns. Eluted protein was loaded into a HiPrep 26/10 Desalting column (Cytiva) to exchange buffer to 20 mM TRIS pH 8.0. Ionic exchange chromatography (IEC) was carried out in a HiTrap CaptoQ ImpRes column (Cytiva) to remove remaining impurities. Purified ACE2 was concentrated in a 50 kDa 4 mL Amicon (Merck), aliquoted and frozen in liquid nitrogen for storage at −80°C.

#### TwinStrep-tagged protein expression and purification

RBD_WT_, RBD_Q498Y_ and RBD_β_baculovirus were used to infect Sf9 cells in a 1:2000 dilution, for 72 hours. Each culture media was collected and recombinant protein was purified from the supernatant using StrepTactin 4Flow® 5 mL cartridge (Iba Lifesciences) and buffers recommended by the manufacturer. Eluted samples were concentrated in 10 kDa 4 mL Amicons (Merck) and further purified by size exclusion chromatography (SEC) using a Superdex 200 10/300 GL column (Cytiva) in a TBS pH 7.4, 1 mM DTT buffer to prevent dimer formation. Protein samples were concentrated using 10 kDa Nanosep columns (Pall Corporation), aliquoted and frozen in liquid nitrogen for storage at −80°C.

### Kinetic characterization using BLI

Affinities of RBD_WT_, RBD_Q498Y_ and RBD_β_ to ACE2 were tested by biolayer interferometry using a BLItz system (Sartorius). His-tagged ACE2 was immobilized onto Ni-NTA coated biosensors (Sartorius) before association of increasing concentrations of each RBD for 120 seconds followed by dissociation using HBS pH 7.4 for 360 seconds. Obtained sensograms were corrected using a blank curve and fitted to a 1:1 Langmuir binding model using BLItz software to characterize binding affinities of each RBD to ACE2.

In another strategy to assess affinities of the different RBDs to ACE2, RBD_WT_was immobilized onto activated amine-reactive sensors. Then, sensors were dipped for 60 seconds into solutions containing 400 nM ACE2, or 400 nM ACE2 pre-incubated with equimolar amounts of RBD_WT_, RBDQ498Y and RBD_β_, prior to dissociation with HBS pH 7.4 for another 60 seconds. Obtained sensograms were corrected using a blank curve.

### RBD_Q498Y_-ACE2 complex crystallization

Sf9 insect cells were infected with RBD_Q498Y_ and TwinStrep-tagged ACE2 (1:2000) separately for 72 hours. Recombinant proteins were purified from each collected supernatant by affinity chromatography using StrepTactin XT 4Flow® 5 mL cartridges (Iba Lifesciences). Eluted RBD_Q498Y_ and ACE2 were then concentrated and buffer-exchanged to TBS pH 7.4 using 10 kDa and 50 kDa 4 mL Amicons (Merck), respectively. Tags were removed by overnight digestion with 3C protease (1:50) at 4°C. 3C-digested proteins were loaded into StrepTactin XT 4Flow® 1 mL gravity flow columns (Iba Lifesciences) and collected in the flow-through for quantification. Equimolar amounts of ACE2 and RBD_Q498Y_ were then pooled together and the formed complex was purified using a Superdex 200 10/300 GL column (Cytiva) and a TBS pH 7.4, 1 mM DTT buffer.

Purified RBD_Q498Y_-ACE2 complex was concentrated to 5.5 mg/mL and screened against different crystallization conditions by the sitting drop vapour diffusion method. Best crystals were obtained in 0.1 M sodium phosphate pH 6.5, 12%w/v PEG 8000, then captured and soaked in mother liquor containing 20% glycerol and cryo-cooled in liquid nitrogen. X-ray diffraction and data collection was performed at BL13-XALOC beamline in the ALBA synchrotron facility.

### X-ray diffraction data processing, structure determination and refinement

Collected data reduction was performed with XDS, then scaled using Aimless in the CCP4i suite excluding 5% of reflections for validation purposes. Structure was solved by molecular replacement using Phaser and previously deposited coordinates for RBD_WT_-ACE2 complex (PDB accession number 6M0J) as template, and refined using Phenix.refine. The final molecule was generated after several cycles of manual building in Coot followed by refinement.

## Accession numbers

Atomic coordinates and structure factors for the RBD_Q498Y_-ACE2 complex have been deposited in the Protein Data Bank under the accession code 7P19.

## Acknowledgements

Jacinto López-Sagaseta is a Ramón y Cajal Investigator. We thank the staff of XALOC beamline at ALBA Synchrotron for their assistance with X-ray diffraction data collection. We thank Maria Gilda Dichiara Rodriguez, Ane Ochoa Echeverria and Adela Rodriguez Fernandez for their excellent technical support.

## Author contributions

Conceptualization: JLS; Methodology: JLS, EEA; Investigation: JLS, EEA; Funding acquisition: JLS; Supervision: JLS; Draft writing: JLS, EEA.

## Supplementary Materials

**Table I.** X-ray data collection and refinement statistics. Statistics for the highest-resolution shell are shown in parentheses.

|  |  |
| --- | --- |
| **Resolution range** | 93.99 - 3.24 (3.35 - 3.24) |
| **Space group** | P 21 21 21 |
| **Unit cell** | 60.591 165.037 228.678 90 90 90 |
| **Total reflections** | 202345 (20961) |
| **Unique reflections** | 37446 (3704) |
| **Multiplicity** | 5.4 (5.7) |
| **Completeness (%)** | 99.43 (99.78) |
| **Mean I/sigma(I)** | 7.87 (1.19) |
| **Wilson B-factor** | 85.19 |
| **R-merge** | 0.2136 (1.508) |
| **R-meas** | 0.2365 (1.661) |
| **R-pim** | 0.09988 (0.6879) |
| **CC1/2** | 0.995 (0.651) |
| **R-work** | 0.245 (0.384) |
| **R-free** | 0.292 (0.408) |
| **RMS(bonds)** | 0.002 |
| **RMS(angles)** | 0.53 |
| **Ramachandran favored (%)** | 95.23 |
| **Ramachandran outliers (%)** | 0.07 |

**Table II.** Atomic contacts between RBD_Q498Y_ and ACE2. Van der Waals, salt bridges and H-bonds are listed based on distance cutoffs of 4 Å, 4.5 Å and 3.4 Å, respectively.

| RBD | ACE2 | Bond type |
| --- | --- | --- |
| Lys417 C<sup>ε</sup> | Asp30 C<sup>γ</sup>, Asp30 O<sup>δ1</sup>, Asp30 O<sup>δ2</sup> | Van der Waals |
| Lys417 N<sup>ζ</sup> | Asp30 C<sup>γ</sup> | Van der Waals |
| Lys417 N<sup>ζ</sup> | Asp30 O<sup>δ1</sup>, Asp30 O<sup>δ2</sup> | Salt bridge |
| Tyr449 C<sup>ε1</sup> | Asp38 C<sup>γ</sup>, Asp38 O<sup>δ1</sup>, Asp38 O<sup>δ2</sup> | Van der Waals |
| Tyr449 C<sup>ζ</sup> | Asp38 C<sup>γ</sup>, Asp38 O<sup>δ1</sup>, Asp38 O<sup>δ2</sup> | Van der Waals |
| Tyr449 OH | Asp38 C<sup>γ</sup> | Van der Waals |
| Tyr449 OH | Asp38 O<sup>δ2</sup>, Asp38 O<sup>δ1</sup> | Hydrogen bond |
| Tyr449 OH | Gln42 C<sup>γ</sup>, Gln42 C<sup>δ</sup>, Gln42 O<sup>ε1</sup> | Van der Waals |
| Tyr449 OH | Gln42 N<sup>ε2</sup> | Hydrogen bond |
| Tyr453 OH | His34 C<sup>γ</sup>, His34 C<sup>δ2</sup>, His34 N<sup>ε2</sup>, His34 C<sup>ε1</sup> | Van der Waals |
| Tyr453 OH | His34 N<sup>δ1</sup> | Hydrogen bond |
| Leu455 C<sup>δ1</sup> | Asp30 O<sup>δ2</sup>, His34 C<sup>δ2</sup>, His34 N<sup>ε2</sup> | Van der Waals |
| Phe456 C<sup>δ1</sup> | Thr27 C<sup>γ2</sup> | Van der Waals |
| Phe456 C<sup>ε1</sup> | Thr27 C<sup>γ2</sup>, Asp30 O<sup>δ2</sup> | Van der Waals |
| Phe456 C<sup>ζ</sup> | Thr27 C<sup>γ2</sup> | Van der Waals |
| Ala475 O | Gln24 C<sup>γ</sup> | Van der Waals |
| Phe486 C<sup>δ1</sup> | Met82 S<sup>δ</sup>, Tyr83 OH | Van der Waals |
| Phe486 C<sup>ε1</sup> | Met82 S<sup>δ</sup>, Met82 C<sup>β</sup>, Tyr83 C<sup>ε1</sup>, Tyr83 C<sup>ζ</sup>, Tyr83 OH | Van der Waals |
| Phe486 C<sup>ζ</sup> | Tyr83 C<sup>ε1</sup> | Van der Waals |
| Asn487 C<sup>γ</sup> | Gln24 O<sup>ε1</sup>, Tyr83 OH | Van der Waals |
| Asn487 O<sup>δ1</sup> | Gln24 O<sup>ε1</sup>, Gln24 C<sup>β</sup>, Gln24 C<sup>γ</sup>, Tyr83 C<sup>ε1</sup>, Tyr83 C<sup>ζ</sup> | Van der Waals |
| Asn487 O<sup>δ1</sup> | Tyr83 OH | Hydrogen bond |
| Asn487 N<sup>δ2</sup> | Gln24 C<sup>δ</sup>, Tyr83 C<sup>ε1</sup>, Tyr83 OH | Van der Waals |
| Asn487 N<sup>δ2</sup> | Gln24 O<sup>ε1</sup> | Hydrogen bond |
| Tyr489 C<sup>δ2</sup> | Lys31 C<sup>γ</sup> | Van der Waals |
| Tyr489 C<sup>ε1</sup> | Thr27 C<sup>γ2</sup> | Van der Waals |
| Tyr489 OH | Phe28 C<sup>β</sup>, Phe28 N, Phe28 C<sup>α</sup> | Van der Waals |
| Tyr489 OH | Tyr83 OH | Hydrogen bond |
| Gln493 C<sup>δ</sup> | Lys31 C<sup>ε</sup>, His34 C<sup>β</sup> | Van der Waals |
| Gln493 O<sup>ε1</sup> | Lys31 C<sup>ε</sup>, His34 C<sup>β</sup>, His34 C<sup>α</sup>, His34 C, Glu35 N, His34 O | Van der Waals |
| Gln493 N<sup>ε2</sup> | Lys31 C<sup>ε</sup>, Lys31 N<sup>ζ</sup> | Van der Waals |
| Gln493 N<sup>ε2</sup> | Glu35 O<sup>ε1</sup> | Hydrogen bond |
| Gly496 C<sup>α</sup> | Lys353 N<sup>ζ</sup> | Van der Waals |
| Gly496 C | Lys353 N<sup>ζ</sup> | Van der Waals |
| Gly496 O | Lys353 C<sup>δ</sup>, Lys353 C<sup>ε</sup> | Van der Waals |
| Gly496 O | Lys353 N<sup>ζ</sup> | Hydrogen bond |
| Tyr498 C<sup>β</sup> | Tyr41 C<sup>ε2</sup> | Van der Waals |
| Tyr498 C<sup>γ</sup> | Tyr41 C<sup>ε2</sup> | Van der Waals |
| Tyr498 C<sup>δ1</sup> | Tyr41 C<sup>δ2</sup>, Tyr41 C<sup>ε2</sup> | Van der Waals |
| Tyr498 C<sup>ε1</sup> | Lys353 N<sup>ζ</sup>, Tyr41 C<sup>δ2</sup>, Tyr41 C<sup>ε2</sup> | Van der Waals |
| Tyr498 C<sup>ε2</sup> | Gln42 O<sup>ε1</sup> | Van der Waals |
| Tyr498 C<sup>ζ</sup> | Gln42 O<sup>ε1</sup> | Van der Waals |
| Tyr498 OH | Asp38 O<sup>δ1</sup>, Gln42 C<sup>γ</sup>, Gln42 C<sup>δ</sup>, Gln42 N<sup>ε2</sup> | Van der Waals |
| Tyr498 OH | Gln42 O<sup>ε1</sup> | Hydrogen bond |
| Thr500 C | Tyr41 OH, Asp355 O<sup>δ2</sup> | Van der Waals |
| Thr500 O | Tyr41 OH, Asp355 C<sup>γ</sup>, Asp355 O<sup>δ1</sup>, Asn330 N<sup>δ2</sup> | Van der Waals |
| Thr500 O | Asp355 O<sup>δ2</sup> | Hydrogen bond |
| Thr500 C<sup>β</sup> | Tyr41 OH, Arg357 NH<sub>2</sub>, Asp355 O<sup>δ2</sup>, Asn330 N<sup>δ2</sup> | Van der Waals |
| Thr500 O<sup>γ1</sup> | Tyr41 C<sup>ζ</sup>, Arg357 NH<sub>2</sub> | Van der Waals |
| Thr500 O<sup>γ1</sup> | Tyr41 OH, Asp355 O<sup>δ2</sup> | Hydrogen bond |
| Thr500 C<sup>γ2</sup> | Arg357 NH<sub>2</sub> | Van der Waals |
| Asn501 N | Tyr41 OH | Van der Waals |
| Asn501 C<sup>α</sup> | Lys353 O, Tyr41 OH | Van der Waals |
| Asn501 C | Lys353 O | Van der Waals |
| Asn501 O<sup>δ1</sup> | Lys353 C<sup>β</sup>, Lys353 C<sup>δ</sup>, Tyr41 C<sup>ε2</sup>, Tyr41 C<sup>ζ</sup> | Van der Waals |
| Asn501 O<sup>δ1</sup> | Tyr41 OH | Hydrogen bond |
| Asn501 N<sup>δ2</sup> | Lys353 C<sup>δ</sup> | Van der Waals |
| Gly502 N | Lys353 C, Gly354 C, Gly354 O | Van der Waals |
| Gly502 N | Lys353 O | Hydrogen bond |
| Gly502 C<sup>α</sup> | Lys353 O, Gly354 C<sup>α</sup>, Gly354 C, Gly354 O | Van der Waals |
| Tyr505 C<sup>β</sup> | Lys353 O | Van der Waals |
| Tyr505 C<sup>γ</sup> | Lys353 C<sup>β</sup>, Lys353 C<sup>α</sup>, Lys353 C, Lys353 O | Van der Waals |
| Tyr505 C<sup>δ2</sup> | Lys353 C<sup>α</sup>, Lys353 C, Lys353 O, Gly354 N | Van der Waals |
| Tyr505 C<sup>ε2</sup> | Lys353 C<sup>α</sup>, Lys353 C, Gly354 N | Van der Waals |
| Tyr505 C<sup>ζ</sup> | Glu37 O<sup>ε2</sup> | Van der Waals |
| Tyr505 OH | Glu37 C<sup>δ</sup>, Glu37 O<sup>ε1</sup>, Glu37 O<sup>ε2</sup> | Van der Waals |

**Figure S1.**
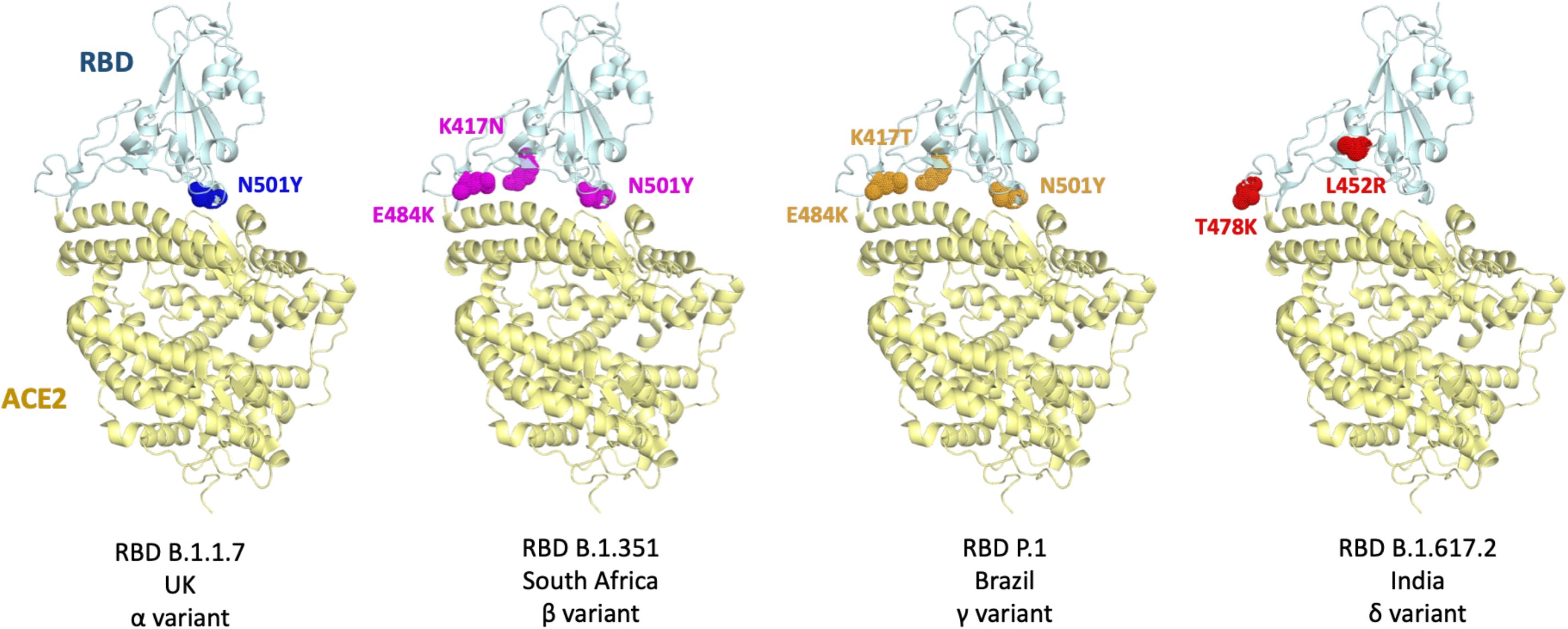
SARS-CoV-2 variants of concern and associated amino acid substitutions in the RBD. Cartoon representation of the RBD_WT_-ACE2 complexes (cyan and pale yellow colors, respectively) using the atomic coordinates deposited in the PDB under the accession number 6M0J. The amino acid substitution for each variant are highlighted with dots and colors.

